# Calorie Restriction Upregulates Islet PD-L1 Signaling and Decreases the Risk of Auto-immune Diabetes Onset in NOD Mice

**DOI:** 10.64898/2026.02.15.705935

**Authors:** Amanda Cambraia, Michael Schleh, Jean-Phillipe Cartailler, Melanie Cutler, Cristiane dos Santos, Gina Many, Hyeyoon Kim, Young-Mo Kim, Ernesto S. Nakayasu, Denis A. Moglienko, Rafael Arrojo e Drigo

## Abstract

Type 1 diabetes (T1D) is an autoimmune disease where beta cells are destroyed by cytotoxic T cells. Calorie restriction (CR) enhances glucose homeostasis and promotes beta cell longevity and was used as therapeutic strategy for T1D prior to the discovery of insulin. However, a significant knowledge gap remains regarding its effects on beta cells during the pathogenesis of autoimmunity. We demonstrate that CR enhances glucose homeostasis, reduces beta cell load, and delays T1D onset in NOD mice. CR induced a largely post-mitotic beta cell state marked by selective loss of beta cell identity markers, reduced DNA damage and beta cell senescence, and increased PD-L1 within the islet microenvironment. This beta cell phenotype correlates with anti-inflammatory and exhausted immune cell states in the NOD islet. Together, these findings indicate that CR improves glucose homeostasis and remodels the islet microenvironment to promote beta cell longevity via a pro-tolerogenic immune microenvironment that reduces the risk for autoimmune diabetes.

## Introduction

Type 1 diabetes (T1D) is the most common chronic autoimmune disease in children and adolescents, and its rising prevalence over the past three decades represents a significant public health concern (*1, 2*). T1D is characterized by immune-mediated destruction of insulin-producing pancreatic beta cells by CD8^+^ cytotoxic T cells and assisted by CD4^+^ T cell, resulting in chronic hyperglycemia, and a lifelong requirement for exogenous insulin treatment (*1, 2*). Both genetic susceptibility and environmental factors contribute to the initiating and persistence of autoreactive immune responses in T1D (*3*). Despite advances in immunotherapy, the complexity and heterogeneity of the T1D onset and progression have so far precluded curative treatments, and the single FDA-approved therapy to delay clinical onset, anti-CD3 antibodies (Teplizumab), extends this period by only 2-3 years (*4-6*).

Beyond genetics and environmental factors, immune activity can be modulated by changes in diet composition, nutrition, and energy intake (*7, 8*). In fact, in 1915 Frederick M. Allen and Elliot P. Joslin promoted “starvation diets” (∼400kcal/day) as a potential treatment for T1D. This approach briefly alleviated hyperglycemia and prolonged survival in some diabetic patients but became rapidly obsolete following the discovery of insulin by Banting and Best (*9*).

More recently, CR has been shown to dampen inflammation and alter the immune cell composition in streptozotocin-treated diabetic mice and in humans with or without autoimmune disease, including multiple sclerosis and rheumatoid arthritis (*7*). Importantly, the impact of CR on immune cell function and their molecular phenotypes can be context-dependent, and it involves (at least in part), reduced mTORC1 activity in immune cells (*7, 10*). Accordingly, mTORC1 inhibition by rapamycin expands CD4+ regulatory T cells (Tregs) and can delay T1D in animals and humans (*11, 12*).

In beta cells, we and others have shown that CR promotes beta cell health, identity, and longevity, while delaying the onset of beta cell aging signatures shared with T1D and type 2 diabetes (T2D) (*13-16*). Mechanistically, CR enhances systemic insulin sensitivity via loss of fat mass loss and prolonged fasting, thereby reducing the secretory burden and reducing the potential for protein misfolding with insulin production in beta cells (*13, 17*). In turn, this promotes a largely post-mitotic phenotype characterized by a significant re-organization of the adult beta cell transcriptome, promoting elevated expression of identity genes and transcription factors, improved energy and proteostasis pathways, strengthened circadian rhythm genes and autophagy, while protecting mitochondria from degradation and suppressing mTORC1 signaling (*13*). This phenotype resembles the concept of beta cell rest, where reduced insulin demand, achieved via exogenous insulin infusion, alleviates the metabolic stress in pre-diabetic beta cells (*18-20*). Approaches that induce a state of beta cell rest or improved glucose homeostasis have been linked to at least some degree of partial preservation of beta cell mass (the “honeymoon effect”) in autoimmune diabetes (*21*). Together, these studies suggest that induction of improved glucose homeostasis, the preservation of endogenous beta cell mass via immunosuppression, and beta cell rest approaches may protect against autoimmune diabetes.

To investigate the impact of CR on T1D onset and progression in a preclinical model, we exposed 8-week-old female NOD mice to ad-libitium (AL) feeding or 20% CR for two months, and performed *in vivo* metabolic phenotyping to assess glucose homeostasis and beta cell function, followed by confocal microscopy, flow cytometry, single nucleus transcriptomics, and cellcell communication analyses to define the molecular changes in islet endocrine cells and islet-associated immune populations. Two months of CR promoted an insulin sensitive state that reduced beta cell insulin secretion and delayed T1D onset in NOD mice. At the tissue level, CR decreased insulitis and reduced the density of immune cells present in active insulitic processes. Moreover, CR promoted a largely postmitotic beta cell state associated with selective loss of beta cell identity markers, reduced DNA damage, and diminished beta cell senescence. Strikingly, CR triggered significant upregulation of the immunomodulatory checkpoint ligand PD-L1 across islet endocrine cells (alpha, beta, and delta). Ligand-receptor analysis mapped islet PD-L1 signaling to PD-1-expressing CD4^+^ Tregs and cytotoxic CD8^+^ T cells, which acquired broad anti-inflammatory and exhausted cell states.

Together, these findings demonstrates that CR can preserves beta cell mass by reshaping beta cell heterogeneity and by activating anti-inflammatory and immunomodulatory pathways that limit immune cell recruitment and effector activation within the islet microenvironment in NOD mice.

## Results

### Calorie restriction (CR) enhances insulin sensitivity to delay diabetes onset in NOD mice

Calorie restriction (CR) can prolong cellular and organismal longevity, including beta cells (*13, 17, 29-31*). These effects are commonly associated with enhanced glucose homeostasis and suppressed pro-inflammatory signaling (*7, 13, 17*). In fact, CR induces a state of reduced demand for beta cell function and insulin release that mimics beta cell rest (*13*), which in turn has been associated with delayed beta cell destruction and preserved beta cell mass in mice and humans with T1D (*19*). We hypothesized that the beneficial effects of CR on systemic and islet-centric glucose homeostasis mechanisms (including beta cells) would delay the onset of T1D in the NOD mouse model. To test this hypothesis, we investigated the impact of 20% CR on NOD mouse glucose homeostasis mechanisms. Here, eight-week-old NOD female mice were exposed to ad-libitum (AL) feeding or 20% CR for 2 months as previously established by us (**Figure 1A**)(*13*). As expected, CR NOD mice were leaner due to significant loss of fat mass relative to lean mass (**Figure 1B-C** and **Supplementary Figure 1A-B**).

**Figure 1.**
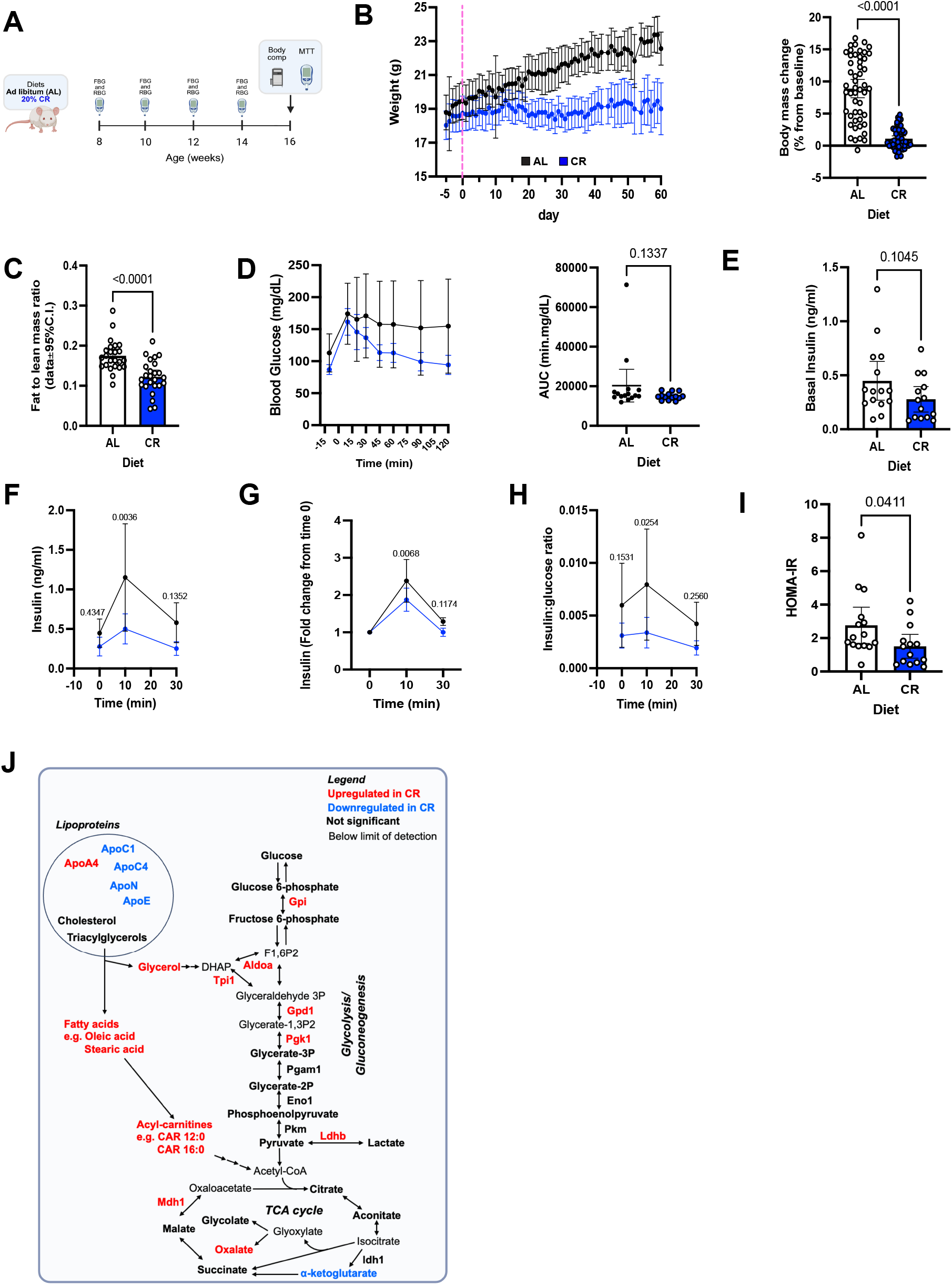
Calorie restriction (CR) modulates beta cell function and delays autoimmune diabetes progression in female NOD mouse. **(A)** Schematic diagram of female NOD mice subjected to ad libitum (AL) or 20% CR for 2 months, starting at 8 weeks old. **(B)** Body weight measured daily for five days before we started the diet until the last day of the regimen and the percent change in body weight relative to baseline. **(C)** Fat-to-lean mass ratio after 2 months on diet. **(D)** Blood glucose levels during the meal tolerance test (MTT) after 2 months on diet and the respective area under the curve (AUC) measurements. **(E)** Basal insulin levels before the MTT, **(F)** Insulin levels during MTT and **(G)** the respective fold change from baseline values. **(H)** Ratio between insulin and glucose values obtained from the MTT. **(I)** HOMA-IR calculated from fasting insulin and glucose values after 2 months on diet. **(J)** Annotated metabolic pathways identified in our metabolomics approaches. Proteins and Metabolites in red and blue are enriched or depleted, respectively, in the blood of CR NOD mice versus AL NOD mice. Molecules in bold black were not significantly altered. In **(D, F, G, and H)**, data were analyzed using a mixed-effects model (REML) with Fisher’s LSD multiple comparisons test, comparing AL vs CR at each time point (**D, F, G** and **H**). For two groups comparison, unpaired two-tailed Student’s t test was performed (**C, D, E** and **I**). Survival was analyzed using Kaplan–Meier survival analysis with log-rank (Mantel–Cox) test. All data presented as mean ±95% confidence intervals (C.l.). In **(B)** *n=* 20 AL and *n=* 20 CR mice per group; In **(C)** *n=* 24 AL and *n=* 25 CR mice per group**; In (D-I)** *n=* 15 AL and *n=* 14 CR mice per group. Panel<u>s **(A,)**and**(J)**</u> were created with BioRender.com.

Next, we assessed glucose homeostasis and beta cell function *in vivo* via an oral mixed-meal tolerance test (MTT), which revealed that CR NOD mice had normal glucose clearance and normal fasting basal insulin levels (**Figure 1D-E**). Notably, CR beta cells secreted significantly less glucose-stimulated insulin than AL mice (**Figure 1F**) and were marked by a lower glucose-dependent beta cell stimulation index (**Figure 1G**). Importantly, despite the relative reduction in circulating insulin levels, the secretory activity of CR beta cells was sufficient to maintain normoglycemia (**Figure 1H**) and likely due to increased peripheral insulin sensitivity (**Figure 1I**). We also performed a multiomics (metabolomics, proteomics and lipidomics) analysis of the blood plasma samples to further study metabolic changes induced by CR (**Figure 1J**). Glucose level remained unaltered, further supporting the findings of the MTT test. None of the glycolysis/gluconeogenesis metabolites had significant changes. However, 6 out of 9 enzymes measured from these pathways were increased in CR, thus suggesting a more efficient carbohydrate metabolism phenotype in CR NOD mice. In terms of lipids, the levels of cholesterol and triacylglycerols remained unaltered. However, the downstream degradation intermediates of triacylglycerols were upregulated, namely glycerol, fatty acids (e.g., oleic and stearic acids), and acylcarnitines (e.g. CAR 12:0 and CAR 16:0). These results indicate that CR NOD mice have increased fatty acid metabolism. Of note, the discrepancy between the maintained levels of triacylglycerols and increased degradation products could be due to shifts in lipoprotein profiles or changes in composition. For instance, we observed an increase in apolipoprotein A4 (APOA4), and decreases in APOC1, APOC4, APOE and APON (**Figure 1J)**.

Together, these findings demonstrate that the metabolic response of female NOD mouse to CR includes significant loss of fat mass, increased lipid metabolism, and dampened nutrient-coupled beta cell insulin secretion and enhanced insulin sensitivity. Surprisingly, CR did not improve glucose clearance during the MTT despite having elevated insulin sensitivity, and which may be explained by elevated hepatic gluconeogenesis levels and impaired hepatocyte insulin signaling in NOD mice (*32*).

### Reduced islet insulitis and enhanced beta cell longevity in CR NOD mice

The progression of T1D is characterized by infiltration of immune cells that target islet beta cells, triggering islet inflammation, and beta cell destruction (*1, 33*). In wild type mice, CR increases peripheral insulin sensitivity to promote a long-lived and post-mitotic state in beta cells associated with enhanced beta cell homeostasis and metabolism (*13*). To determine if CR has similar effects on NOD beta cell biology and function, we monitored the progression of spontaneous diabetes in AL and CR mice during fed and fasted states throughout the dietary intervention. To determine the timeline of autoimmune diabetes onset in our colony, we performed blood glucose measurements and mice with consecutive random blood glucose (RBG) of at least 250mg/dL and fasting blood glucose (FBG) of at least 150mg/dL were classified as diabetic (**Supplementary Figure 2A**). Importantly, at least 50% of female NOD mice in our facility develops spontaneous diabetes within 19 weeks of age, which is well within the range of our experimental timeline (**Supplementary Figure 2B**). Here, we observed that 2 months of 20% CR had a significant effect on reducing the onset rates of spontaneous diabetes in CR NOD mice versus AL-fed mice in both fasting and fed conditions (**Figure 2A and Supplementary Figure 2C**).

**Figure 2.**
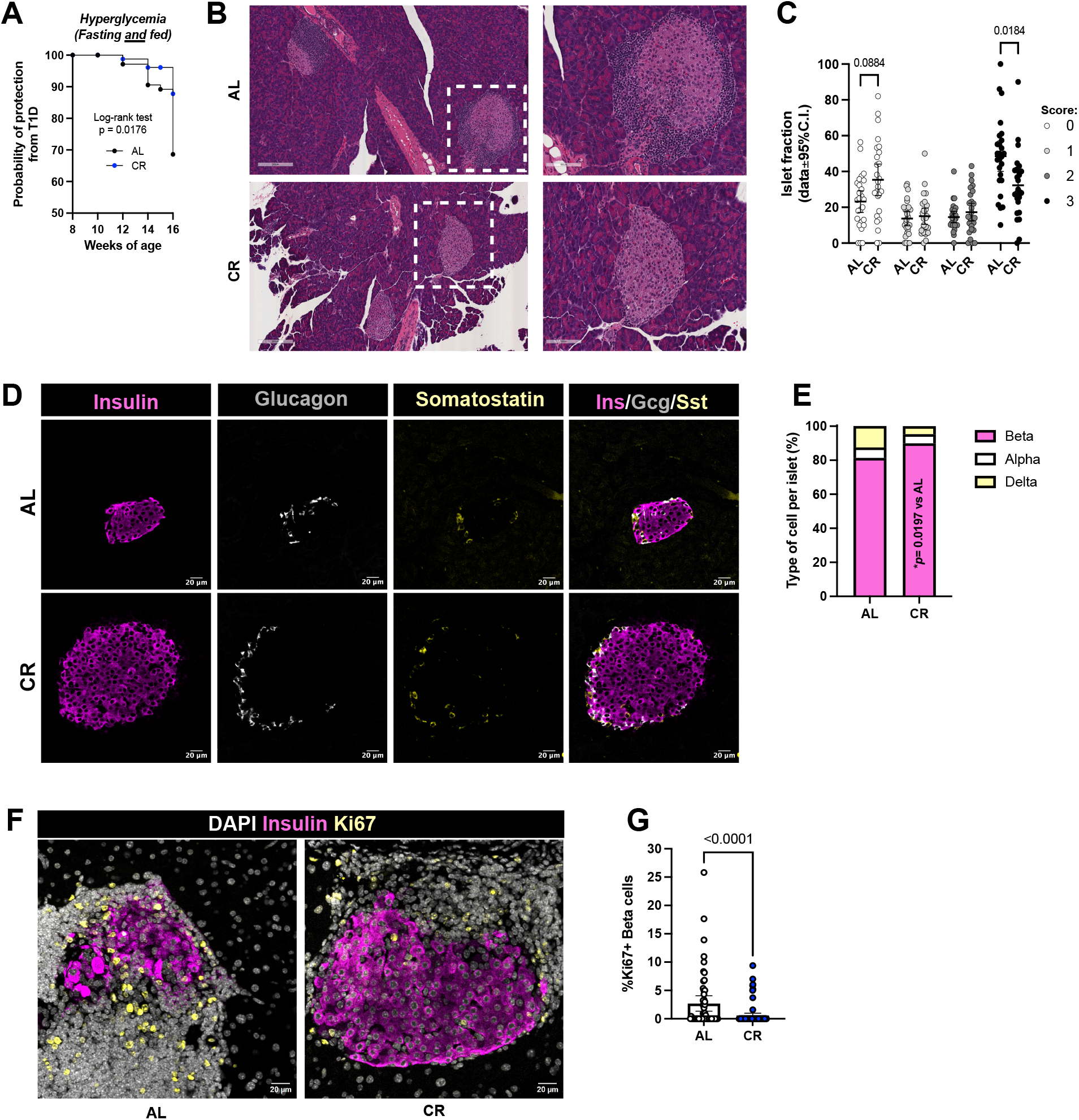
CR delays hyperglycemia onset and preserves pancreatic islet architecture. **(A)** Kaplan-Meier analysis showing probability rates of hyperglycemia onset in NOD female mice after 2 months on diet, assessed by fasting and fed blood glucose measurements. **(B)** Representative hematoxylin and eosin (H&E) stained pancreatic sections from AL and CR mice after 2-months on diet showing islet morphology and immune infiltration. Dashed boxes indicate regions shown at higher magnification. **(C)** Quantification of insulitis severity score per islet in AL and CR groups. **(D)** Representative islets from AL and CR mice after 2 months on diet stained with Insulin (magenta), Glucagon (White) and Somatostatin (yellow). **(E)** Quantification of each cell type per diet group, expressed as percentage. Scale bar = 20 μm. **(F)** Representative islets from AL and CR mice after 2 months on diet stained with DAPI (white), Insulin (magenta) and Ki67 (yellow) and **(G)** quantification of Ki67 positive beta cells per diet group. Scale bar = 20 μm. Data are presented as mean with 95% confidence interval (C.I.) and student’s t-test was used for two groups comparisons. Hyperglycemia was defined as two with consecutive RBG ≥ 250mg/dL and FBG ≥ 150mg/dL. Survival was analyzed using Kaplan–Meier survival analysis with log-rank (Mantel–Cox) test (**A**). Data were analyzed using two-way repeated-measures ANOVA, with Sidak’s post-hoc test (**C**). For two groups comparison, unpaired two-tailed Student’s t test was performed (**G**). All data presented as mean ±95% confidence intervals (C.l.). In **(A)**, data pooled from *n=*4 different cohorts, with a total of 60 AL and 55 CR mice at the start of the diet intervention; in **(C)** *n=* 26 AL and *n=* 27 CR mice per group; in **(E)** *n=* **34** AL islets and *n=* 65 CR islets; and in *n=* 4 mice per group(**G)** *n=* 35 AL islets and *n=* 26 CR islets; and in *n=* 5 mice per diet group.

Next, to determine if this reduction in T1D onset was due to decreased autoimmune attack to CR NOD islets, we dissected pancreases from AL and CR mice after 2 months on diet and analyzed islet structure and inflammation using immunohistochemistry (IHC). Analysis of H&E-stained pancreases followed by islet scoring revealed that CR NOD mice have more islets without immune cell infiltration (i.e., insulitis) and significantly less islets with the most severe levels of immune cell infiltration (**Figure 2B-C**). Moreover, imaging of NOD islet cytoarchitecture identified that CR NOD mice have higher beta cell mass than AL mice (**Figure 2D-E**), thus suggesting that CR preserves NOD beta cells. Next, to determine if this phenotype was explained by changes in beta cell proliferation rates, we quantified beta cell proliferation in AL and CR islets using Ki67 immunostaining. This revealed a significant reduction in beta cell proliferation rates in CR islets (**Figure 2F-G**), thus indicating that the elevated beta cell mass in CR NOD mice is likely explained by preservation of existing beta cells instead of the formation of new beta cells.

### Single nucleus RNA sequencing reveals dissociation of beta cell longevity and identity signatures in CR NOD islets

We have previously shown that functional and molecular adaptations of beta cells to CR involve a significant re-organization of the beta cell transcriptome that promotes beta cell identity and homeostasis mechanisms (*13*). Therefore, we performed single nucleus RNA sequencing (snRNA-seq, 10X genomics) of nuclei extracted from isolated AL and CR NOD islets to determine the impact of CR on the molecular signature of endocrine cells and immune cells within the islet microenvironment (**Figure 3A**). After quality control and filtering, we analyzed a total of 45,051 single nuclei (AL= ∼17051 and CR= ∼28000) to identify all major islet cell types, vascular and accessory cell types, and several immune cell types (**Figure 3B**).

**Figure 3.**
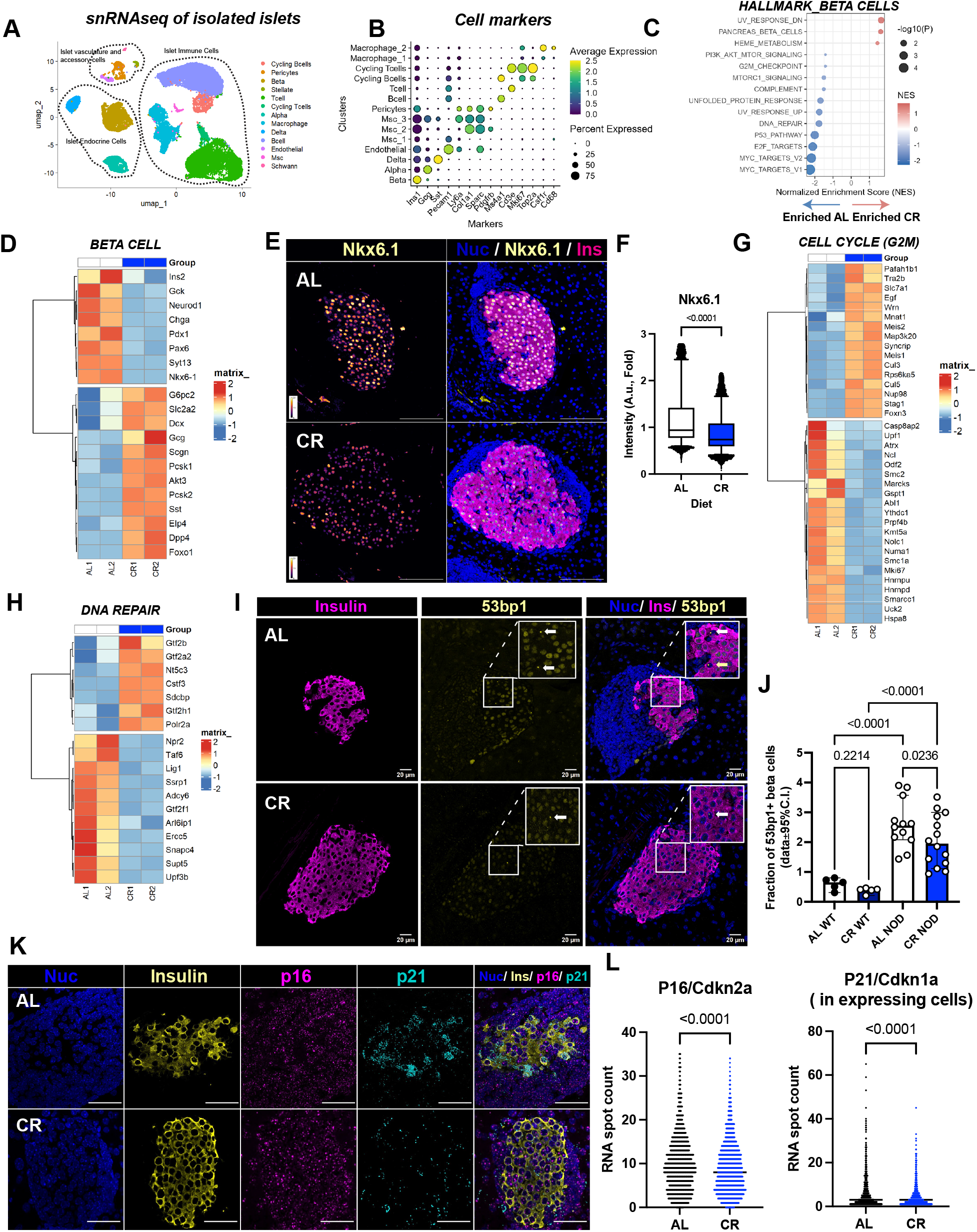
Single-cell RNA sequencing of pancreatic islets reveals beta cell identity remodeling in female NOD mice. **(A)** Uniform Manifold Approximation and Projection (UMAP) of the integrate single-nucleus RNA-seq (snRNA-seq) data obtained from isolated pancreatic islets from AL and CR NOD female mice after 2 months on diet. **(B)** Dot plot showing expression of selected marker genes across clusters used for cell type annotation. Dot size represents the percentage of nuclei expressing each gene, and color intensity indicates average normalized expression. **(C)** Dot plot showing significantly enriched hallmark gene sets in beta cells, with the normalized enrichment score (NES) plotted on the x-axis. Dot size represents −log10(P value), and color indicates NES, with red denoting positive enrichment and blue denoting negative enrichment for CR beta cells. **(D)** GSEA hallmarks for gene sets Hallmark_PANCRE-ATIC_BETA_CELLS. **(E)** Representative images of islets from AL and CR mice after 2 months on diet stained with DAPI (blue), NKX6.1 (yellow) and Insulin (magenta). Left panels show Nkx6.1 signal alone, and right panels show merged images. **(F)** Quantification of NKX6.1 fluorescence intensity is shown on the right. **(G)** GSEA hallmarks for gene sets Hallmark_G2M. **(H)** GSEA hallmarks for gene sets Hallmark_DNA_REPAIR. **(I)** Representative islets from AL and CR mice after 2 months on diet. Slides were stained with DAPI (blue), insulin (magenta) and 53BP1 (yellow). **(J)** Quantification shows the fraction of 53bp1+ beta cells per islet in wild-type (WT) and NOD mice under AL or CR conditions. Each dot represents one islet, and bars indicate 95% confidence intervals (CI). 53BP1+ beta cells were defined as insulin-positive cells containing one or more discrete nuclear 53BP1 foci. **(K)** Representative RNAscope images of pancreatic islets from AL and CR mice after 2-months on diet stained for DAPI (Blue), Insulin (Yellow), p16 (Magenta) and p21 (Cyan). **(L)** Quantification of RNAscope signal showing RNA spot counts per beta cell for p16/Cdkn2a (left) and p21/Cdkn1a (right, in expressing cells only) in AL and CR groups. Each dot represents one beta cell. For two groups comparison, unpaired two-tailed Student’s t test was performed (**F** and **L**). Data were analyzed with one-way ANOVA with Benjamini, Krieger and Yekutieli’s post-hoc test (**J**). All data presented as mean ±95% confidence intervals (C.l.). In **(E)** *n=* 6 AL and 6 CR mice per diet group; In **(I)** *n=* 5 AL wild-type mice, *n=* 5 CR wild-type mice, *n=* 12 AL NOD mice and *n=* 14 CR NOD mice; In **(K)** *n=* 6 AL and *n=* 6 CR mice per diet group.

Next, we performed pseudobulk analysis in subset cell populations to identify differentially expressed genes followed by Gene Set Enrichment Analysis (GSEA) to identify pathways modulated by CR in NOD beta cells (**Supplementary Table 1**). We found that CR differentially regulated pathways associated with DNA damage and repair, cell proliferation, mTORC1 signaling, unfolded protein response (UPR), beta cell identity, and circadian rhythm (**Figure 3C, Supplementary Figure 3 and Supplementary Table 1**). Specifically, CR upregulated beta cell function genes *Glut2/Slc2a2, G6pc2, Pcsk1/2, Ero1b*, and circadian regulators *Arnt/Bmal, Tef*, and *Hlf* (**Supplementary Table 1)** – all of which are known to be responsive to CR in wild type beta cells (*13*). Of note, CR beta cells have elevated expression of alpha cell (*Gcg, Tm4sf4*) and delta cell (*Sst*) lineage genes, as well as reduced expression of major transcription factors that regulate beta cell identity (i.e., *NeuroD1, Pdx1, Nkx6-1, Ins2* and *Pax6*) and of insulin secretion machinery genes (i.e., *Chga, Chgb, Glp1r, Gipr, Syt7*) (**Figure 3D, Supplementary Table 1**). This data strongly suggests that while CR promotes the expression of glucose metabolism and circadian rhythm genes in NOD beta cells, these cells also have impaired beta cell identity features. To validate this hypothesis *in situ*, we quantified NKX6-1 in AL and CR NOD islets by IHC, revealing a significant reduction in beta cell Nkx6-1 protein levels in CR NOD islets, thus supporting our initial snRNA-seq observations (**Figure 3E-F**).

In T1D, loss of beta cell identity and function are thought to be caused by chronic (and unresolved) ER stress due to significant cytotoxic stress signaling from the autoimmune system and that contributes to beta cell death (*34, 35*). Given that CR NOD islets contain more beta cells that are less proliferative (**Figure 2E-G**), we investigated whether this could be explained by reduced ER stress and downstream UPR activation. GSEA pathways analysis for UPR activation in CR NOD beta cells included higher expression of the main ER stress sensors *Atf6, Ire1/Ern1*, and *Eif2ak3/Perk* in CR beta cells (**Supplementary Figure 3B, Supplementary Table 1**). However, CR beta cells do not appear to have an active ER stress signaling response given that the expression of ER stress responsive genes downstream of *Atf6/Perk/Ire1a* signaling (i.e., *Hspa5/Bip* and *Ddit3/Chop*) and other ER stress response proteins (e.g., *Bag3, Ddit4, Pdia5/6, Calr, Atf3*) is reduced in CR islets (**Supplementary Figure 3B**). Moreover, CR NOD beta cells have significant downregulation of ER-associated degradation (ERAD) pathway genes, and no significant upregulation of pro-apoptotic genes was observed (**Supplementary Table 1**). In fact, CR NOD beta cells have upregulation of genes linked to ER protein folding (*Bag2, Dnacj1, Sil1*), protein maturation (*Stt3a, Tusc3*), protein translocation and export (*Sec13, Sec24b/d, Tram1*), and ER redox proteins (*Ero1b, Man1a/Man1a2*). Together, this response suggests that these cells have increased capacity for ER proteostasis to mediate misfolding potential.

The elevated expression of DNA damage response and repair genes in AL mice suggests that CR NOD beta cells might accumulate lower DNA damage and senescence (like wild type CR beta cells (*14*)). Consistent with this interpretation, GSEA pathway analysis in CR NOD beta cells revealed enrichment of DNA repair and cell cycle-associated mechanisms (**Figure 3G-H**), including upregulation of *Wrn* gene, a key regulator of genome stability and replication stress and telomere maintenance (*36*) (**Figure 3G**). In addition, CR beta cells exhibited upregulation of core RNA polymerase II machinery components, including *Polr2a, Gtf2b* and *Gtf2a2* and mRNA processing and nucleotide metabolism genes (i.e., *Cstf3* and *Nt5c3a*) (*37*) (**Figure 3H**). This coordinated enhancement of transcriptional and post-transcriptional mechanisms, rather than activation of cell proliferation (**Figure 2F-G and Figure 3G**), indicates a protective state of transcriptional integrity and enhances cellular resilience to stress. To validate these findings, we immunostained AL and CR NOD pancreases with anti-53BP1 antibodies to quantify DNA damage accumulation, which revealed a significant decrease in the fraction of 53bp1-positive beta cells in CR NOD mice (**Figure 3I-J**). Of note, CR NOD beta cells had higher 53bp1 positivity than age-matched wild type CR mice (**Figure 3I-J**). Next, we used the RNAscope platform to perform *in situ* mRNA hybridization for the beta cell senescence markers *Cdkn1a/p21* and *Cdkn2a/p16* and observed decreased levels of both senescence markers in CR NOD beta cells (**Figure 3K-L**), thus indicating that CR beta cells have reduced senescence signatures.

Together, this data demonstrates that CR NOD beta cells do not experience significant levels of ER stress or apoptosis; instead, their increased survival in CR NOD mice (**Figures 1-2**) correlates with a relatively immature and largely post-mitotic beta cells state with reduced accumulation of DNA damage and cellular senescence, consistent with reduced expression of Ki67+ cell in CR NOD mice.

### CR dampens islet immune cell pro-inflammatory signatures

Infiltration of T cells into pancreatic islets is a hallmark of T1D linked to islet inflammation, breakdown of beta cell immune tolerance, and beta cell death (*35*). In other autoimmune disease models, CR has been shown to exert anti-inflammatory effects that can alter disease progression (*7*). Here, led by the observation that CR NOD mice have decreased levels of insulitis, we hypothesized that CR could be influencing the phenotype of the immune cell system surrounding the islet to promote beta cell longevity and health.

To test this hypothesis, we first quantified the relative cell density in the insulitis process to find that CR NOD islets have significantly fewer immune cells in their periphery than AL mice (**Supplementary Figure 4A**). Additional IHC and confocal microscopy experiments using anti-ki67 antibodies revealed that lower cell density in the insulitic process is due to (at least in part) lower rates of cell proliferation (**Supplementary Figure 4B**). Next, we performed pancreas IHC using anti-CD4 and -CD8 antibodies to investigate the density of specific T cell subpopulations in and around AL and CR NOD islets. Here, we found that CR specifically increases the density of intra-islet CD4+ cells (**Figure 4A-B**). To further investigate these observations, we performed flow cytometry analysis of immune cells isolated from the draining pancreatic lymph node (pLN) of AL and CR NOD mice after 2 months on diet. We analyzed CD4- and FOXP3-expressing cells among CD45^+^CD3e^+^ T cells to identify FOXP3^+^ regulatory T (Treg) cells (**Figure 4C and Supplementary Figure 4D**). This analysis identified a significant increase in frequencies of CD4^+^ FOXP3^+^ T cells in CR NOD pancreases, thus suggesting that CR could increase the proportion of immunoregulatory CD4+ Treg cells to decrease T-cell mediated inflammation and apoptosis and thereby facilitate increased in NOD beta cell survival during CR (**Figure 4D**).

**Figure 4.**
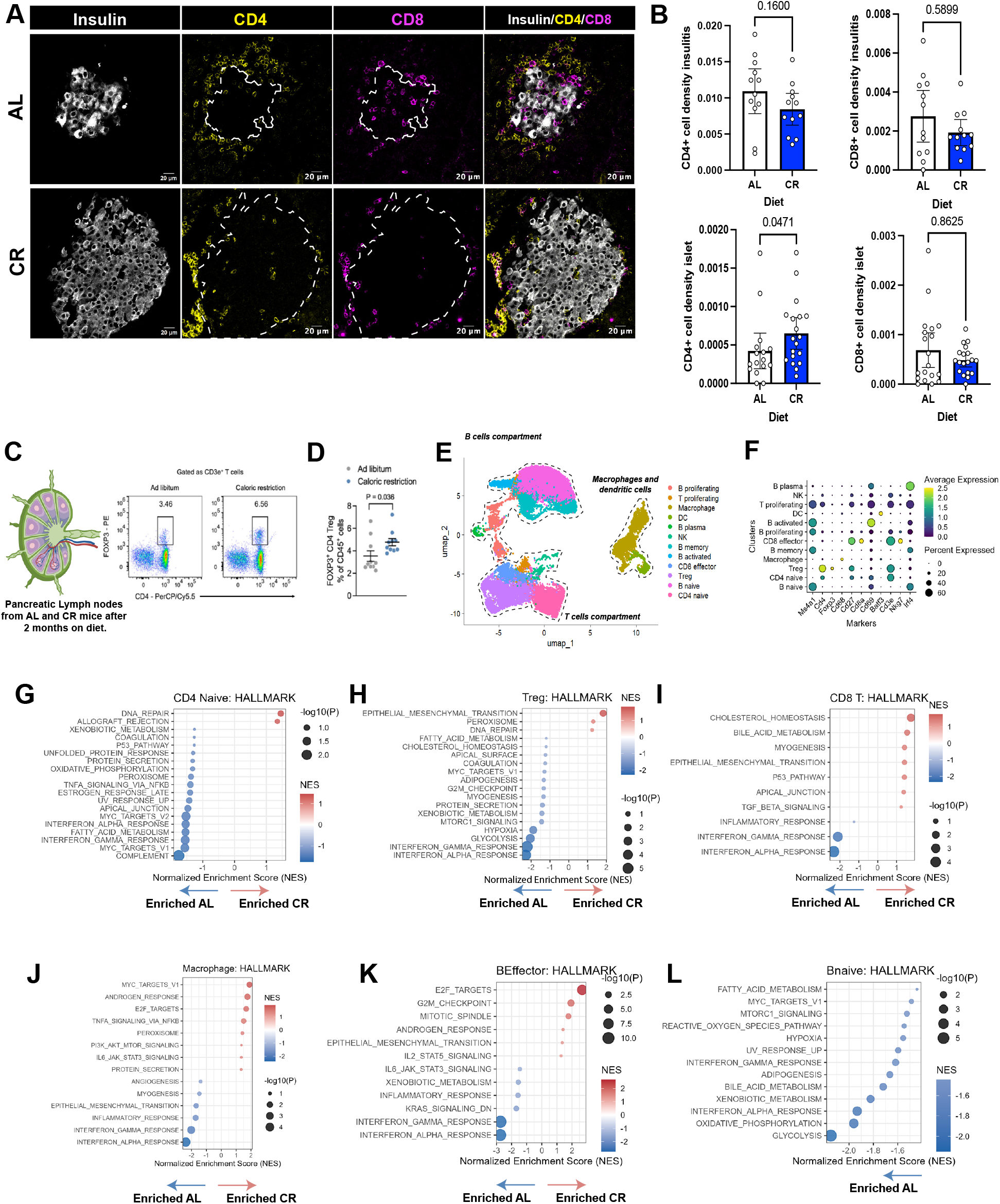
CR modulates islet immune composition and immune cell functional states. **(A)** Representative islets from AL and CR mice after 2 months on diet stained with Insulin (white), CD8 (yellow) and CD4 (magenta). Scale bar: 20um. **(B)** Density of CD4+ and CD8+ T cells within the islet and within the insulitic process in AL and CR female NOD mice. Bars indicate 95% CI. **(C)** Schematic of pancreatic lymph node (PLN). **(D)** Quantification of Treg frequency (FOXP3^+^ of CD4^+^ T cells) in pLNs from AL and CR mice (n=10 mice per group; two-sided unpaired t-test). **(E)** UMAP projection of immune cells from snRNA-seq of isolated islets. **(F)** Dot plot with average expression of immune cells markers. **(G)** GSEA of CD4+ naïve T cells comparing CR and AL conditions. Pathways with positive NES are upregulated in CR, whereas pathways with negative NES are downregulated. Only significantly enriched pathways are shown. **(H)** GSEA of CD4+ Treg cells comparing CR and AL conditions. Pathways with positive NES are upregulated in CR, whereas pathways with negative NES are downregulated. Only significantly enriched pathways are shown. **(I)** GSEA of CD8+ T cells comparing CR and AL conditions. Pathways with positive NES are upregulated in CR, whereas pathways with negative NES are downregulated. Only significantly enriched pathways are shown. **(J)** GSEA of Macrophages comparing CR and AL conditions. Pathways with positive NES are upregulated in CR, whereas pathways with negative NES are downregulated. Only significantly enriched pathways are shown. **(K)** GSEA of Effector B cells comparing CR and AL conditions. Pathways with positive NES are upregulated in CR, whereas pathways with negative NES are downregulated. Only significantly enriched pathways are shown. **(L)** GSEA of Naïve B cells comparing CR and AL conditions. Pathways with positive NES are upregulated in CR, whereas pathways with negative NES are downregulated. Only significantly enriched pathways (FDR q < 0.05) are shown. For two groups comparison, unpaired two-tailed Student’s t test was performed (A, B, D and F). All data presented as mean ±95% confidence intervals (C.l.). In **(B)** *n=* 12 islets per diet group were analyzed for CD4+ and CD8+ T cell density in the insulitis, *n=* 19 islets per diet group were analyzed for CD8+ T cell density in the islets and *n=* 16 AL and 19 CR islets were analyzed for CD4+ T cell density in the islets. (**E)** was created with BioRender.com.

To gain a broader view of immune cell populations in AL and CR NOD islets, we applied snRNAseq analysis to dissect the impact of CR on specific immune cell subpopulations found within the islet microenvironment (**Figure 4E**). Using this approach, we performed high resolution sub-clustering of the islet immune cell populations (shown in Figure 3A) to identify proliferating T cells (*CD3e*^*+*^ *Mki67*^*+*^), CD8 Effector T cells (CD8T) (*CD3e*^*+*^ *CD8a*^*+*^ *CD27*^*+*^), regulatory CD4 T cells (Tregs) (*CD3e*^*+*^ *high CD4*^*+*^ *CD27*^*+*^ *Foxp3*^*+*^), naïve CD4 T cells (*low CD4*^*+*^ and *Foxp3 negative*), macrophages (*CD3e negative, CD68*^*+*^), dendritic cells (DCs) (*CD3e negative, Batf3*^*+*^), natural killer cells (NKs) (*CD3e*^*+*^ *Nkg7*^*+*^), and distinct subpopulations of B cells, including plasma (*Ms4a1 negative, CD69*^*+*^ *Irf4*^*+*^), naïve (*Ms4a1*^*+*^ *CD69*^*+*^ *Irf4*^*+*^), activated (*Ms4a1*^*+*^ *high CD69*^*+*^ *low Irf4*^*+*^), and proliferating (*Ms4a1*^*+*^ *Mki67*^*+*^) B cells (**Figure 4F**). We also observed a trend towards an increase in the relative density of CD4 Tregs and a reduction in naïve CD4 cells in CR NOD islets (**Supplementary Figure 4C**). Next, we performed GSEA analysis to define how CR impacts their molecular phenotypes. Strikingly, CR promoted significant downregulation in pro-inflammatory signaling across all major immune cell subpopulations identified, including CD4 Tregs, naïve CD4s, CD8Ts, macrophages, and B cell compartments (**Figure 4G-**L). Specifically, this signature was largely characterized by downregulation of interferon (alpha and gamma) responses, as well as by changes in tumor necrosis factor alpha (TNFα) signaling, lipid and carbohydrate metabolism pathways, and cell cycle regulators in naïve CD4s, CD4 Tregs, CD8T, and B cell subpopulations (**Figure 4H-M and Supplementary Table 2**).

Together, these results demonstrate that CR reshapes the immune cell landscape in NOD islets by dampening the density and broad suppression of pro-inflammatory signaling in innate and adaptive immune cells.

### Increased beta cell PD-L1 in CR islets correlates with CD4 and CD8T cell exhaustion signatures

Our observations that CR promotes reduced insulitis and increased FOXP3^+^ Tregs in the pLN, as well as anti-inflammatory immune cell phenotypes, and preserved beta cell mass, led us to hypothesize that CR could activate immune checkpoint signaling between beta cells and immune cells to mitigate autoimmune aggression.

To test this hypothesis, we searched for the expression of immunomodulatory genes in CR beta cells in our snRNAseq dataset. We found that the critical immune checkpoint molecule *CD274*/*PD-L1* was among the top genes positively modulated by CR (**Figure 5A, Supplementary Table 1**). We validated these findings using IHC and confocal microscopy and found a 30-50% increase in PD-L1 protein levels in beta cells *in situ* **(Figure 5B, Supplementary Figure 5B)**. Elevated *Cd274/Pd-l1* gene expression was also found in CR alpha cells and delta cells (**Supplementary Table 3-4)**, thus suggesting that regulation of Cd274/Pd-l1 expression by CR is not a beta cell specific event. Next, to determine the targets of cell-cell communication networks involving *Cd274/Pd-l1* signaling in AL and CR NOD islets, we applied the cell-cell communication inference algorithm CellChat (*28*) to analyze our islet snRNAseq data. This approach identified that elevated PD-L1 signaling from CR islet endocrine cells (i.e., alpha, beta, and delta cells) targeted cytotoxic CD8T and CD4 Tregs, and that this communication was absent in AL islets (**Figure 5C**). This data suggests that CR enhances islet–driven PD-L1 axis that targets CD4 and CD8 compartments.

**Figure 5.**
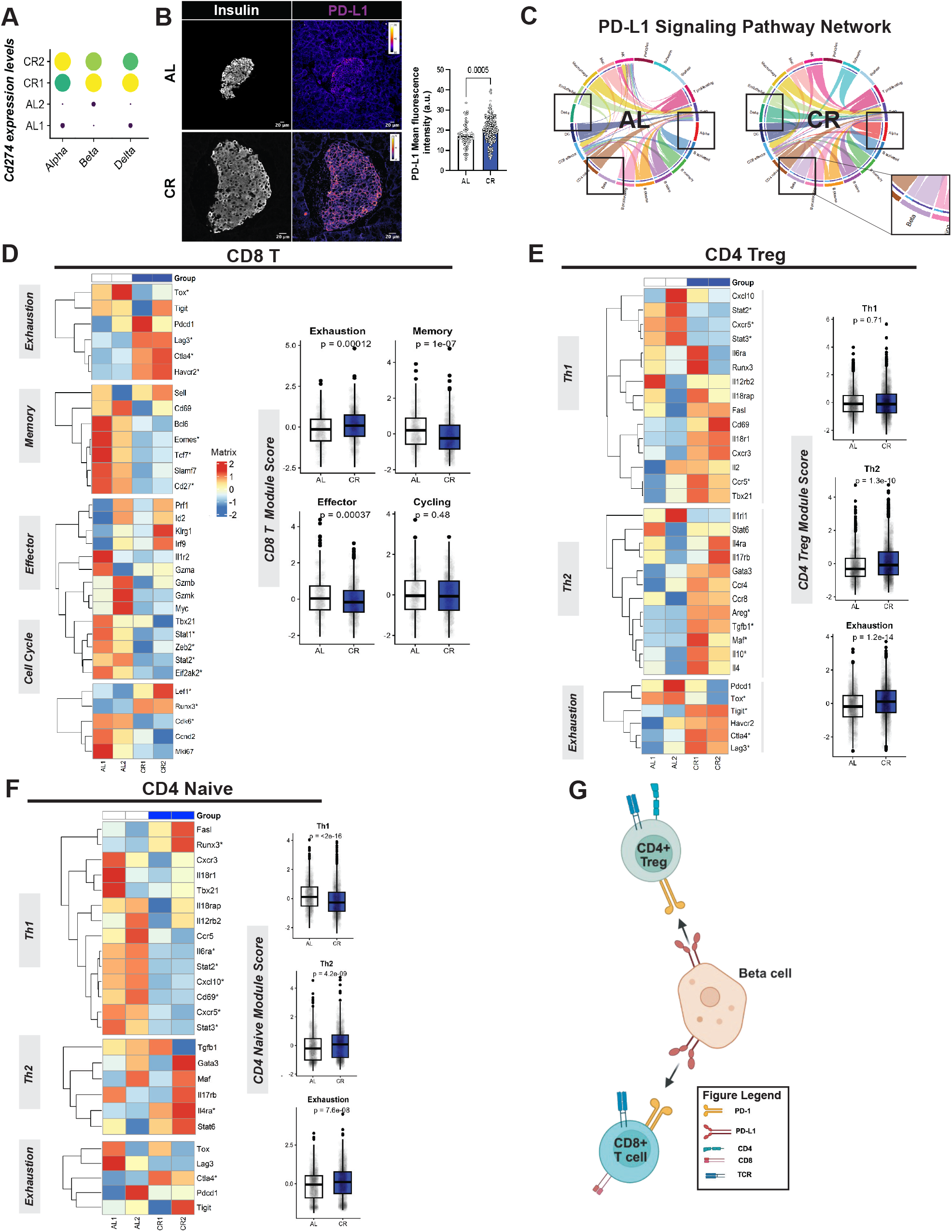
CR modulates PD-L1 signaling and T-cell states in pancreatic islets of female NOD mice. **(A)** Dot plot showing Cd274 (PD-L1) expression levels across islet endocrine cell types (alpha, beta, and delta cells) in AL and CR female NOD mice. **(B)** Representative islets from AL and CR mice showing PD-L1 expression. White dotted lines delineate islet boundaries. Right, quantification of PD-L1 mean fluorescence intensity within islets. **(C)** PD-L1 signaling pathway networks inferred from single-cell RNA-seq data, shown separately for AL and CR conditions. Chord diagrams illustrate predicted ligand–receptor interactions involving PD-L1 between endocrine and immune cell populations. Highlighted regions indicate interactions altered by caloric restriction. **(D)** Heatmap of gene expression signatures associated with CD8+ T-cell functional states (exhaustion, memory, effector, and cell cycle) in AL and CR mice. Right, module score quantification for each functional state. Data are shown as box-and-whisker plots; statistical analysis was performed using a two-tailed Mann–Whitney test. **(E)** Heatmap of gene expression signatures defining CD4+ naïve T-cell states, including Th1, Th2, and exhaustion programs, across AL and CR conditions. Right, corresponding module score comparisons between groups. **(F)** Heatmap of transcriptional programs associated with CD4+ regulatory T cells (Tregs), highlighting Th1-like, Th2-like, and exhaustion-related signatures. Right, module score quantification comparing AL and CR groups. **(G)** Schematic model summarizing the proposed mechanism whereby calorie restriction enhances PD-L1–mediated signaling in islet endocrine cell types, contributing to altered CD8+ T-cell exhaustion and CD4+ T-cell regulatory states, ultimately promoting islet immune tolerance. Gene list used for Th1 module score: *Tbx21, Runx3, Cxcr3, Cxcr5, Ccr5, Il2rb2, Il18r1, Il18rap, Fasl, Cxcl10, Cd69, Il2, Il6ra, Stat2* and *Stat3*. Gene list used for Th2 module score: *Gata3, Il4, Il5, Il13, Stat6, Il4ra, Ccr4, Ccr8, Il1rl1, Il17rb, Areg, Maf, Tgfb1* and *Il10*. Genes used for Exhaustion module score: *Pdcd1, Lag3, Tigit, Ctla4, Tox* and *Havcr2*. For two groups comparison, unpaired two-tailed Student’s t test was performed (A, B, D and F). All data presented as mean ±95% confidence intervals (C.l.). **(B)** *n=* 8 AL and *n=* 11 CR mice per diet group. **(G)** was created with BioRender.com.

The PD-1/PD-L1 pathway contributes to the preservation of local immune tolerance within pancreatic tissue and draining lymph nodes, potentially by inducing T cell exhaustion and functional suppression of effector activation (*38*). In addition, NOD mice lacking PD-1 or PD-L1 develop accelerated type 1 diabetes (*38-40*). Moreover, alternative pathways can induce T cell exhaustion, namely Galectins (via *Tim3*), CD86 (via *Ctla4*), and *Fgl1* (via *Lag3*) (*41-43*). Notably, in CR NOD mice, islet endocrine cells were the major cell types inferred to be communicating with T cells via PD-L1, whereas macrophages and endothelial cells were linked to Cd86 and galectin signaling, respectively (**Supplementary Figure 5C and D**). Therefore, we hypothesized that this elevated “pro-exhaustion” tone from CR islet cells would lead to elevated T cell exhaustion signatures and help explain the observed suppression of pro-inflammatory pathways in CD4+ and CD8+ T cell subpopulations and linked to beta cell survival in CR mice (**Figure 4H**). To address this, we created a panel with well-established CD4 or CD8 T cell exhaustion gene markers (**Supplemental Table 5**) and determined an “exhaustion module score” for each major T lymphocytes subpopulations cell identified in our snRNAseq. Indeed, we observed significantly elevated exhaustion scores in CR CD8+ NOD T cells that resulted in attenuation of memory and effector phenotypes (**Figure 5D**). Consistent effects were observed in naïve CD4+ T cells, where CR increased exhaustion and Th2 phenotypes, while decreasing Th1. (**Figure 5E**) Similarly, Tregs increased exhaustion and Th2 phenotypes upon CR, and notably upregulated anti-inflammatory cytokines Il10 and Tgbf1.

Together, these findings suggest that CR significantly increases islet endocrine cell PD-L1 levels that likely limit effector activation from CD4+ and CD8+ cell types, which adopt an exhausted molecular profile consistent with suppression of cytotoxic autoimmunity (**Figure 5H**).

## Discussion

In this study, we investigated the impact of calorie restriction (CR) on glucose homeostasis mechanisms and on beta cell health and identity in NOD mice. Here, we applied a combination of *in vivo* phenotyping, single nucleus transcriptomics and bioinformatics, and confocal microscopy to establish the relationship between CR-dependent expression of PD-L1 in islet endocrine cells and an increase in immune cell exhaustion signatures with increased beta cell survival and lower risk of autoimmune diabetes in NOD mice. In contrast to our previous CR studies in adult WT mice (*13*), CR did not improve overall glucose clearance and failed to induce beta cell rest despite a significant loss of body fat mass, increased lipid metabolism, and elevated peripheral insulin sensitivity. This may be explained by an impairment of beta cells to secrete enough insulin in response to rising glucose levels (Figure 1) as well as elevated gluconeogenesis flux in NOD mice (*32*) (Figure 1). Importantly, and like WT beta cells, CR NOD beta cells are largely post-mitotic; however, they have limited expression of beta cell identity and function factors (i.e., *Nkx6-1, Pdx1, Chga*). Therefore, CR NOD beta cells are relatively immature, and which explains their insulin secretion phenotype *in vivo* (Figures 2-3). Importantly, CR NOD beta cells can sustain normoglycemia and survive for longer periods of time versus AL mice.

These results support the link between CR and CR-associated increases in peripheral insulin sensitivity with negative regulation of beta cell turnover and enhanced beta cell longevity. In fact, this data suggests that a more mature beta cell identity is not required to achieve and sustain beta cell longevity during autoimmune diabetes in mice. Instead, beta cell longevity in CR NOD mice is characterized by lower levels of DNA damage and senescence accumulation, as well as upregulation of the immuno-modulatory molecule PD-L1 (also found in CR alpha cells and CR delta cells). While we did not detect downregulation PERK signaling, several genes associated with the mTORC1 pathway were downregulated in CR NOD beta cells and that could contribute to elevated PD-L1 levels (*44, 45*). Moreover, this PD-L1-high and immature beta cell phenotype is consistent with previous observations in NOD mice where immature beta cells resist immunological attack via increased expression PD-L1 or inactivation of ER stress sensors (*34, 46-48*). Together, this data suggests that CR promotes a shift in beta cell heterogeneity towards a more resilient and immature beta-cell population that is sufficient to support normoglycemia when insulin sensitivity is optimal.

The significant increase in PD-L1 expression in CR NOD beta cells suggested that beta cell survival could be due to immuno-suppression of effector T cells in the islet periphery. This idea is supported by different observations. First, analysis of CD45+ cells in the pLN revealed a higher number of FOXP3^+^ regulatory T cells in CR NOD mice, thus suggesting that CR can increase the density of regulatory T cells in a critical organ involved in the priming and expansion of islet-reactive T cells. Previous effects in Treg density and function have been observed in NODs and in humans with T1D, where they contribute to impaired peripheral tolerance mechanisms (*49-53*)). Therefore, the enrichment of pLN FOXP3^+^ CD4 T cells may limit the activation and the pathogenic differentiation of autoreactive T cells prior to their migration into the CR pancreas. In addition, CR also promoted expression of tolerogenic signaling pathways within the NOD islet microenvironment. Here, cell–cell communication network analysis revealed significant remodeling of T cell exhaustion in CD4 and CD8 T cells, namely from galectins (from endothelial cells), PD-L1 (from major islet cells), and CD80 (from macrophages) in response to CR feeding. Accordingly, this pro-exhaustion cell-cell communication network is correlated with a broad anti-inflammatory and exhausted phenotype of effector CD8+ T and CD4+ T cell populations (Figure 5), and which can explain the longer survival of CR beta cells.

Overall, our findings support a model where CR remodels the progression of autoimmune diabetes through coordinated metabolic, cellular, and immunological adaptations. CR enhances beta-cell resilience by reducing cellular senescence and identity while promoting immune checkpoint signaling. CR shifts the phenotype of immune regulation at both lymphoid priming sites and within the islet, where major endocrine cells have increased PD-L1 levels that likely causes T-cell exhaustion via PD-1/PD-L1–dependent communication.

### Limitations of this study

Our study demonstrates the beneficial effects of CR on promoting adult NOD mouse beta cell health, which triggers a selective loss of identity and reduces beta cell senescence. This phenotype is associated with elevated PD-L1 expression in islet endocrine cells and associated with a broad anti-inflammatory and exhausted cell state phenotype in CD4+ and CD8+ T cell populations. This data is consistent with previous CR studies in autoimmune disease models, including multiple sclerosis (*7*). However, our study did not investigate the role of the PD-L1 receptor PD-1 in the response of CR NOD mice, nor did we expand our analysis to non-endocrine (stromal, endothelial) and professional antigen presenting cells (e.g., B cells, macrophages) to investigate their potential role in CR NOD mouse phenotype. These are important studies that will be carried out in the future.

## Acknowledgements

We are thankful to the outstanding team at the Vanderbilt Mouse Metabolic Phenotyping Center (MMPC, RRID: SCR_021939) for all the assistance with *in vivo* glucose homeostasis tests (supported by NIH grants DK135073, DK020593). Insulin was measured by the Vanderbilt Analytical Services Core (DK020593). This research was also supported by recruitment funds from the Vanderbilt’s Department of Molecular Physiology and Biophysics and NIH grant 1R01DK138141 to RAeD, and NIH R01AI192909 to D.A.M.

## Author contributions

A.C, M.S., C.D.S. and R.A.D designed experiments. A.C., M.C., G.M., H.K., Y-M. K., E.S.N, C.D.S, D.A.M., R.A.D. collected data and performed data analysis and interpretation. A.C and R.A.D wrote the manuscript.

## Supplementary Figure Legends

**Supplementary Figure 1**: (**A**) Lean mass and (**B**) fat mass. Data are presented as mean with 95% confidence interval (C.I.). A n = 24 AL mice and 25 CR mice. For two groups comparison, unpaired two-tailed Student’s t test was performed (**A** and **B**). All data presented as mean ±95% confidence intervals (C.l.). In (**A-B)** *n=* 15 AL and *n=* 15 CR mice per group.

**Supplementary Figure 2: (A)** Blood glucose levels in NOD female mice under AL or CR over 2 months of dietary intervention, measured in fed and fasted states. **(B)** Probability of hyperglycemia onset over time in female NOD mice housed in the Vanderbilt animal facility. **(C)** Probability of hyperglycemia onset over time in fed or fasted female NOD mice under AL or CR over 2 months of dietary intervention. Survival was analyzed using Kaplan–Meier survival analysis with log-rank (Mantel–Cox) test (A-C). All data presented as mean ±95% confidence intervals (C.l.). In **(A-C)**, data pooled from *n=*4 different cohorts, with a total of 59 AL and 53 CR mice at the start of the diet intervention.

**Supplementary Figure 3: (A)** GSEA hallmarks for gene sets Hallmark_UNFOLDED_PRO-TEIN_RESPONSE. **(B)** GSEA hallmarks for gene sets Hallmark**_**MYC_TARGETS_V1. **(C)** GSEA hallmarks for gene sets Hallmark_MTORC1_SIGNALING. **(D)** GSEA hallmarks for gene sets Hallmark_COMPLEMENT. **(E)** GSEA hallmarks for gene sets Hallmark_P53_PATHWAY. **(F)** GSEA hallmarks for gene sets Hallmark_E2F_TARGETS.

**Supplementary Figure 4: (A)** Quantification of insulitis area normalized to islet area in pancreatic sections from AL and CR female NOD mice after 2 months on diet. Each dot represents one islet. **(B)** Quantification of Ki67+ cells within the insulitis in AL and CR groups. Each dot represents one islet. **(C)** Relative distribution of T-cell populations in AL and CR female NOD mice, shown as stacked bar plots. T cells were classified into CD4 naïve, regulatory T cells (Treg), CD8 effector, proliferating T cells, and natural killer (NK) cells based on transcriptomic markers. Bar graphs depict the frequency of each population within total CD4^+^ or CD8^+^ T cells, as indicated.. **D)** Flow cytometry gating strategy scheme.

**Supplementary Figure 5: (A)** Chord diagrams depict inferred cell–cell communication through the CD86 co-stimulatory pathway in islets from AL and CR female NOD mice. Edge width reflects relative interaction strength, and colors indicate the sending cell population. **(B)** Chord diagrams illustrate galectin-mediated communication networks in AL and CR islets. Cell–cell communication was inferred using single-nucleus RNA-sequencing data and network analysis.

## Supplementary Table Legends

**Supplementary Table 1** – List of genes identified by GSEA analysis of AL and CR beta cells.

**Supplementary Table 2 -** List of genes identified by GSEA analysis of islet immune cell types in AL and CR mice.

**Supplementary Table 3** – List of differently expressed genes in alpha cells.

**Supplementary Table 4** – List of differently expressed genes in delta cells.

**Supplementary Table 5 –** List of exhaustion T cell markers for CD4 and CD8 cell types.

