## Supplementary Figures for "Calorie Restriction Upregulates Islet PD-L1 Signaling and Decreases the Risk of Auto-immune Diabetes Onset in NOD Mice"

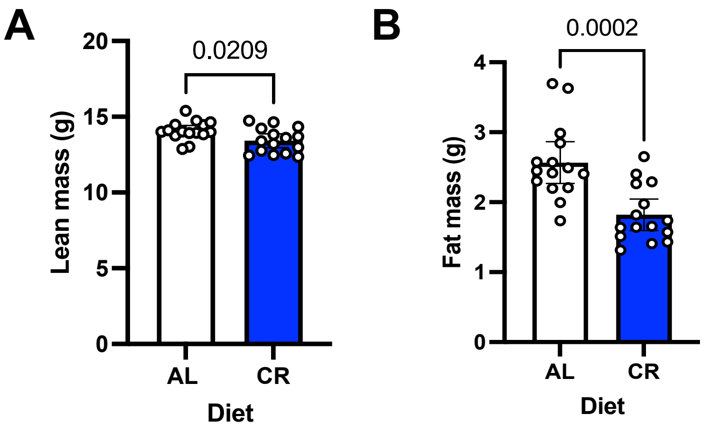


**Supplementary Figure 1**

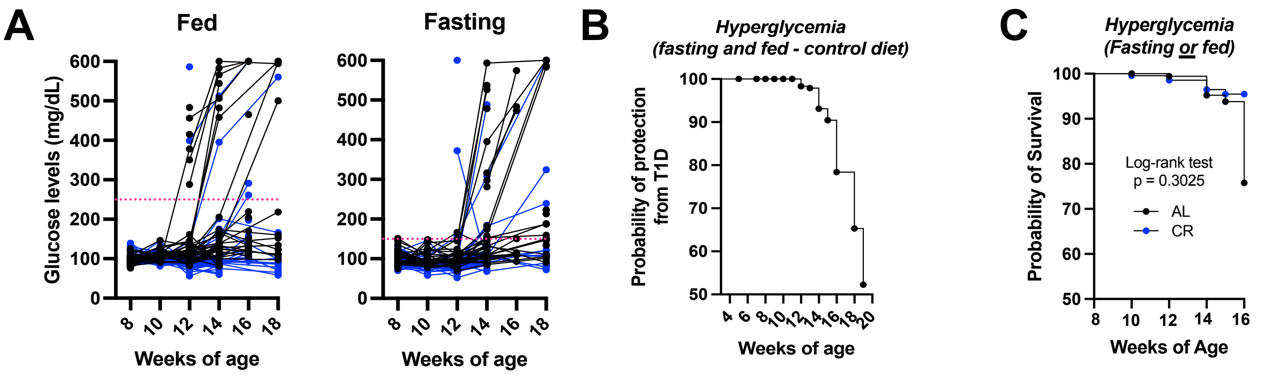


**Supplementary Figure 2**

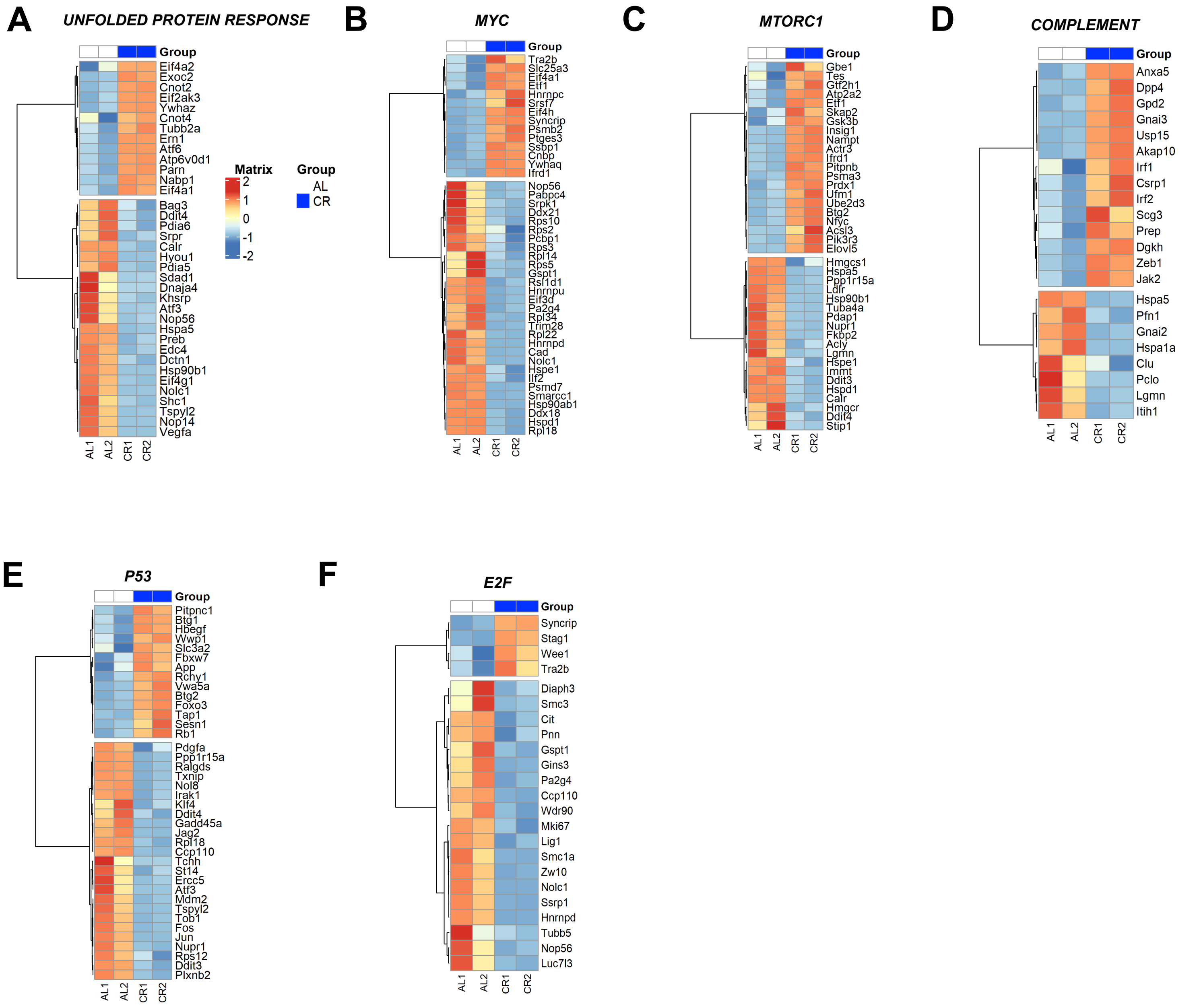


**Supplementary Figure 3**

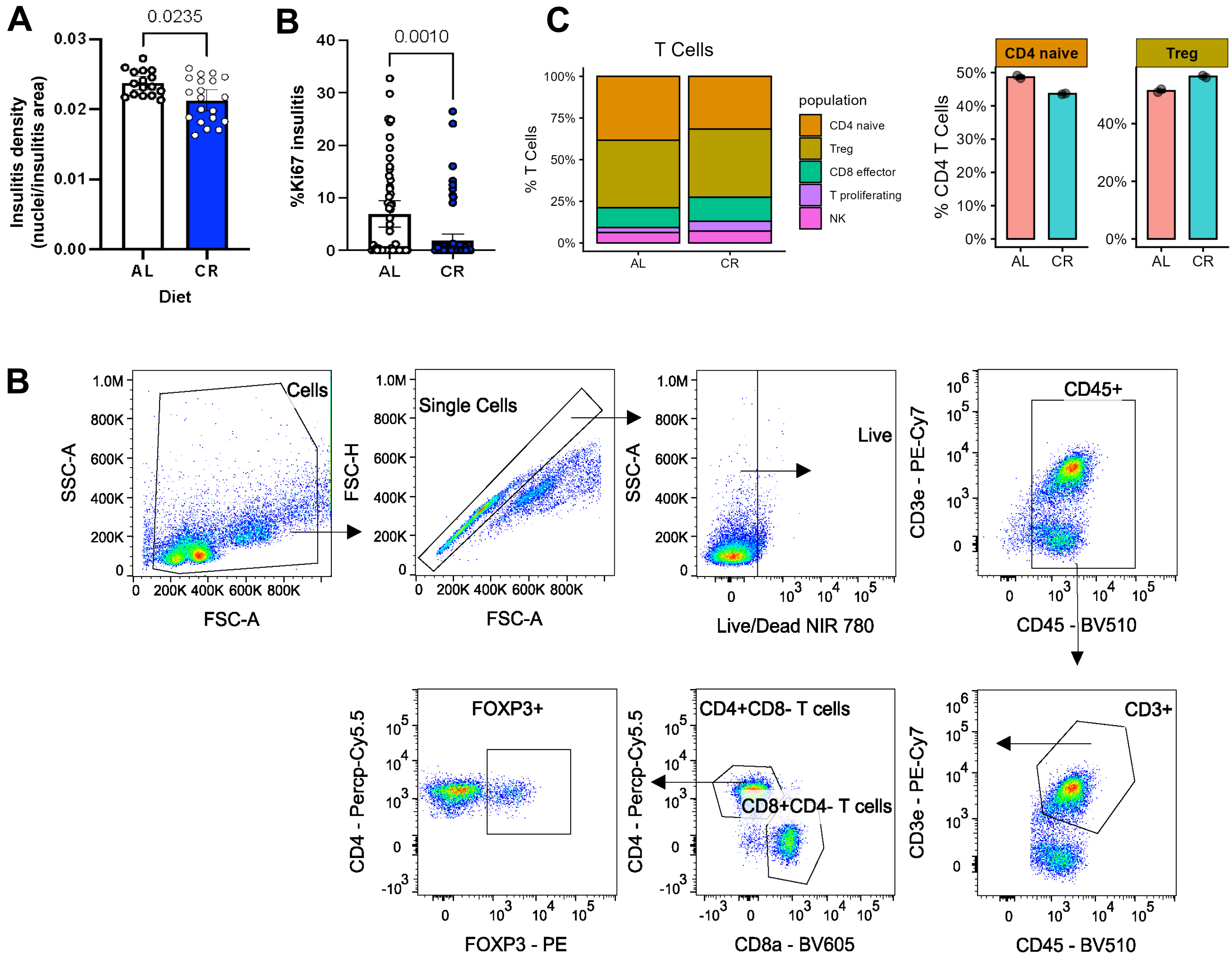


**Supplementary Figure 4**

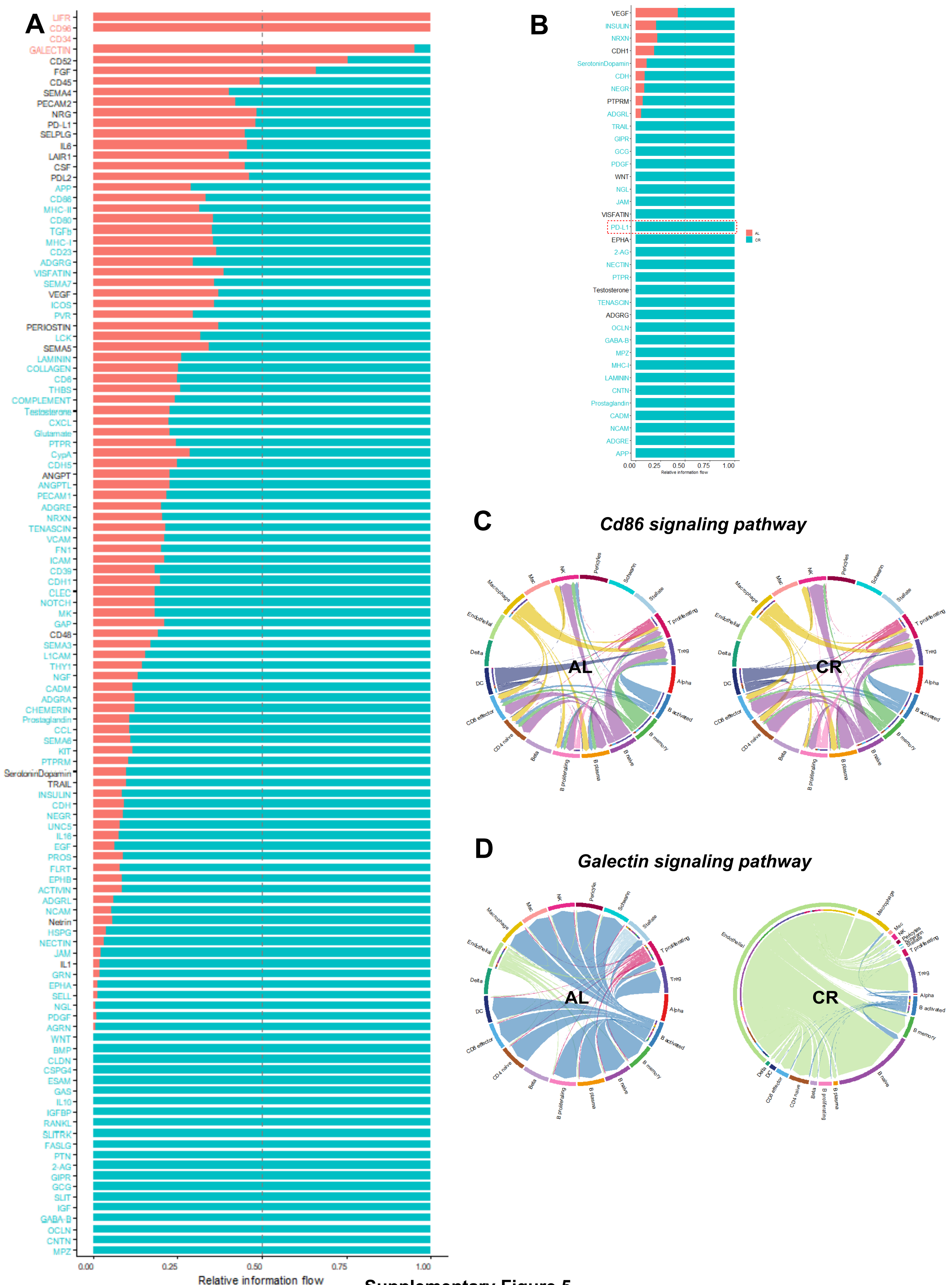


Supplementary Figure 5
